# Comprehensive monitoring of tissue composition using in vivo imaging of cell nuclei and deep learning

**DOI:** 10.1101/2022.10.03.510670

**Authors:** Amrita Das Gupta, Jennifer John, Livia Asan, Carlo Beretta, Thomas Kuner, Johannes Knabbe

**Affiliations:** Department of Functional Neuroanatomy, Institute for Anatomy and Cell Biology, Heidelberg University, Germany; Department of General Psychiatry, Centre for Psychosocial Medicine, Heidelberg University, Germany; Current address: Department of Neurology and Center for Translational Neuro- and Behavioral Sciences (C-TNBS), University Hospital Essen, Germany

## Abstract

Comprehensive analysis of tissue composition has so far been limited to ex-vivo approaches. Here, we introduce NuCLear (<u>Nu</u>cleus-instructed tissue <u>c</u>omposition using deep <u>lear</u>ning), an approach combining in vivo two-photon imaging of histone 2B-eGFP-labeled cell nuclei with subsequent deep learning-based identification of cell types from structural features of the respective cell nuclei. Using NuCLear, we were able to classify almost all cells per imaging volume in the secondary motor cortex of the mouse brain (0.25 mm^3^ containing ∼25000 cells) and to identify their position in 3D space in a non-invasive manner using only a single label throughout multiple imaging sessions. Twelve weeks after baseline, cell numbers did not change yet astrocytic nuclei significantly decreased in size. NuCLear opens a window to study changes in relative abundance and location of different cell types in the brains of individual mice over extended time periods, enabling comprehensive studies of changes in cellular composition in physiological and pathophysiological conditions.

## Introduction

Understanding the plasticity and interaction of different brain cell types in vivo has been a limiting factor for the investigation of large-scale structural brain alterations associated with diverse physiological and pathophysiological states. Until now, to investigate the effect of an experimental intervention on multiple cortical cell types and their respective spatial interactions, each would require an individual labelling strategy using a set of specific promoters and fluorophores. This would require prioritizing cell types from the outset and thereby limit the scope to a small subset of the population. With multiple cell types to consider, the number of experiments and animals investigated may become intractable. To date, studies assessing whole tissue composition quantitatively have primarily employed ex vivo approaches using manual or automated analysis of antibody staining in tissue sections (Ero, Gewaltig et al. 2018), isotropic fractionation (Herculano-Houzel and Lent 2005), in situ hybridization (Nagendran, Riordan et al. 2018) or single cell sequencing (Zeisel, Hochgerner et al. 2018). Other studies used manual (Schmitz and Hof 2007) or automated stereological approaches (Zhang, Yan et al. 2017, Pakkenberg, Olesen et al. 2019). All these studies were conducted ex vivo and a subset required the dissolution of the studied organ’s cellular architecture. The ability to identify multiple cell types and their location in 3D space within a single subject, here referred to as ‘composition’, in the integrity of the living brain to quantify and observe changes over time delivers new opportunities.

Here, we propose an experimental approach to study neurons and glial cells, especially, astroglia, microglia and oligodendroglia, as well as endothelial cells in the same mouse in vivo, using a single genetically modified mouse line expressing a fusion protein of histone 2B and enhanced green fluorescent protein (eGFP) in all cell nuclei. Our approach uses a deep learning method, implementing artificial neural networks to classify each nucleus belonging to a specific cell type, making it possible to perform a <u>nu</u>cleus-instructed tissue <u>c</u>omposition analysis using deep <u>lear</u>ning (NuCLear). Additionally, determining the precise coordinates of every nucleus within the imaged space allows one to assess spatial relations of cells, e.g. the degree of cell clustering vs. even distribution of cells. This can be utilized as an indirect marker of glial territory size, which has been proven to be relevant in pathogenesis of disease such as Alzheimer’s (Bouvier, Jones et al. 2016), or give information about neuron-glia-vasculature proximity (Zisis, Keller et al. 2021). Apart from the application in longitudinal in vivo imaging of the mouse brain, the concept of this technique could be applicable to many other cellular imaging techniques such as confocal and widefield imaging of different organs, when ground truth data for classifier training is available. This will make NuCLear a powerful approach for future analyses of large-scale automated tissue composition.

## Results

For the comprehensive identification and tracking of cell type composition in the mouse brain over the course of weeks and even months, a mouse line constitutively expressing a human histone 2B-eGFP (H2B-eGFP) fusion protein in all cell nuclei was imaged using 2-photon microscopy after implantation of a chronic cranial window (Figure 1A, Supplementary Figure 1A). Volumetric images taken from this mouse line show a nuclei distribution resembling the well-known DAPI staining (Supplementary Figure 1B, C). The idea behind the approach was to use nuclei as proxy for cells and train a neuronal network with ground truth data to classify nuclei belonging to distinct cell types. In brief, the proposed method consists of three main steps: (1) Automated nuclei segmentation in the raw data (detection and labelling of nuclei), (2) feature extraction and (3) classification of individual nuclei using pretrained classifiers for each desired cell type (Figure 1B, C, C(i)). In our study we selected five different cell types for classification: neurons, astroglia, microglia, oligodendroglia, and endothelial cells. Neurons were further classified into inhibitory or excitatory subtypes.

**Figure 1.**
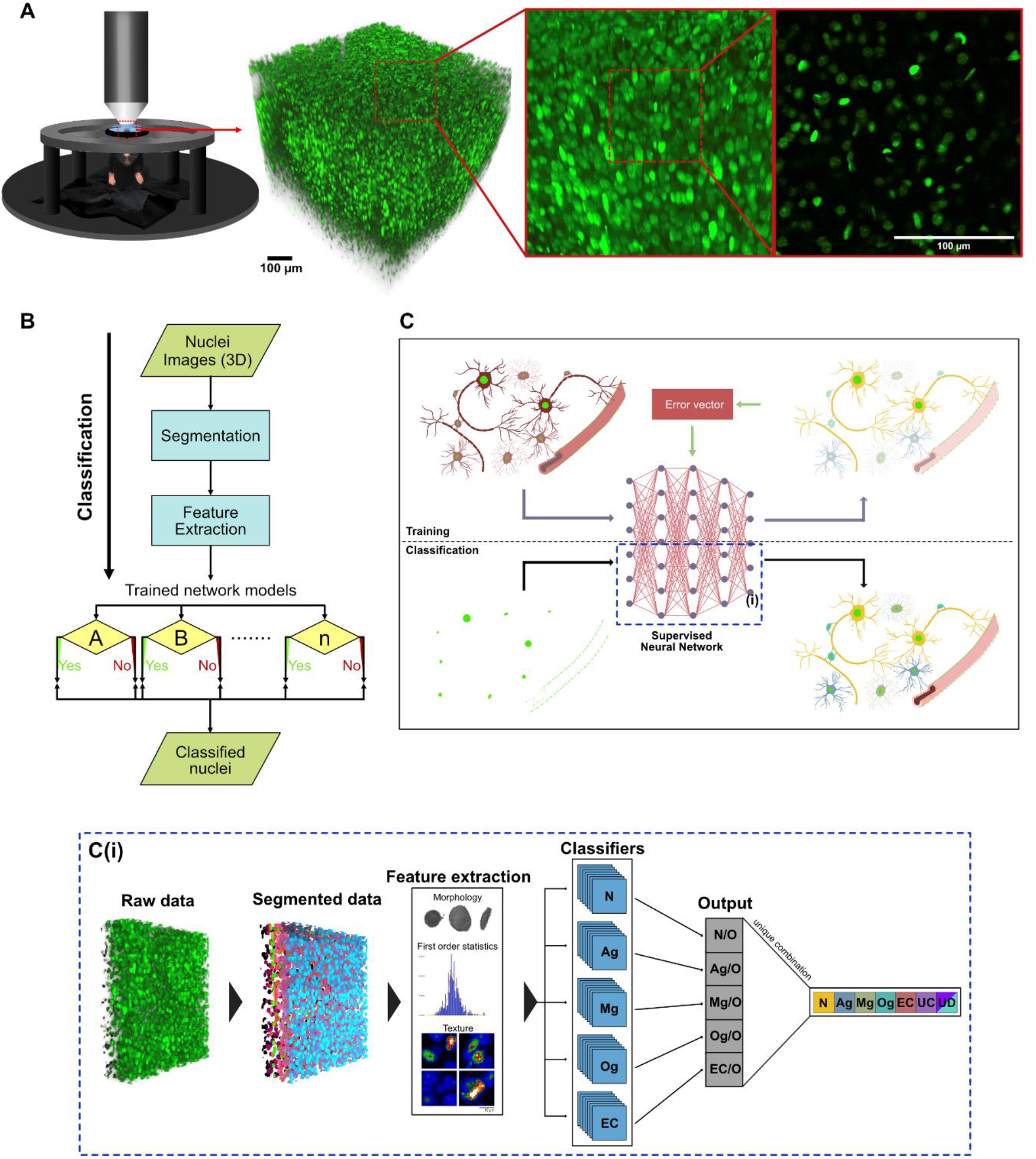
Imaging of cell nuclei and classification into different types. **(A)** Illustration showing acquisition of a 700 µm x 700 µm x 700 µm image volume of cell nuclei in the neocortex using in-vivo 2-photon (2P) microscopy in a histone 2B-eGFP (H2B-eGFP) mouse. 3D reconstruction and single imaging plane. **(B)** Flow diagram showing the classification process from acquisition of the 2P images until the final classification. **(C)** Illustration visualizing a simplified process of the training and classification algorithms. Using an overlay of red (only one cell type labelled per mouse) and green (nuclei with histone 2B-egfp expression: bright green) fluorescent images, ground truth data were obtained to train a supervised neural network to classify nuclei into cell types. Yellow: Neurons, blue: Astroglia, green: Microglia, turquoise: Oligodendroglia, red: Endothelial cells **(C(i))** Visualization of the classification process described in (B) and the blue dashed area in (C). Image volumes were obtained from H2B-eGFP mice, each nucleus was automatically segmented. Features were extracted and each nucleus was classified using pretrained classifiers. (N = Neuron, Ag = Astroglia, Mg = Microglia, Og = Oligodendroglia, EC = Endothelial cells, UC = unclassified cells, UD: undecided cells (classified as belonging to multiple classes)).

To be able to train a classifier for each cell type, ground truth data had to be generated. In addition to the H2B-eGFP-labeled nuclei, a red fluorescent marker protein was introduced (tdTomato, DyLight 549 or mCherry) to mark a specific cell type. The colocalization of both the red and green label allowed us to assign each nucleus to a specific cell type and to extract ground truth data for nucleus classification in the green channel. Nuclei belonging to a specific cell type were manually selected and their features were used to train a neuronal network classifier in a supervised way (Figure 2A). To avoid a bleed through effect of the red fluorescent protein tdTomato into the eGFP channel, which could influence the quality of the classification, we induced the Cre-ERT2-dependent expression of tdTomato via intraperitoneal injection of tamoxifen (Supplementary Figure 1D, right) for cell type identification *after* the first imaging of the nuclei (Figure 2B, overlapping emission spectra for eGFP and tdTomato are shown in Supplementary Figure 1E, bleed through of the tdTomato signal into the eGFP channel is demonstrated in Supplementary Figure 1F). As microglia were the only cells to change their positions after induction with tamoxifen, we chose acute injection of tomato-lectin coupled to DyLight 594 to label these cells, which did not produce any signal alterations in the eGFP-channel (Supplementary Figure 1G). Neurons were labelled using a cortically injected AAV expressing the mCherry red fluorescent protein under the neuron-specific synapsin promotor (syn1-mCherry), which did not affect the eGFP-signal due to its specific fluorescence properties with a more right-shifted emission spectrum compared to tdTomato (Supplementary Figure 1E, G). To further distinguish between different neuronal subtypes, excitatory and inhibitory neurons were labelled by AAV-mediated expression of CamKIIα-mCherry and mDLX-mRuby2 respectively (Supplementary Figure 2A, B). To prepare the ground truth datasets of tdTomato expressing mice, nuclei were manually selected from the pre-induction timepoint, wherever the green and red fluorescence signals overlapped in the post-induction timepoint images (Figure 2C, Supplementary Figure 4B). For the microglial and neuronal datasets, nuclei that overlapped in the green and red channel were manually selected (Figure 2C, Supplementary Figure 2B).

**Figure 2.**
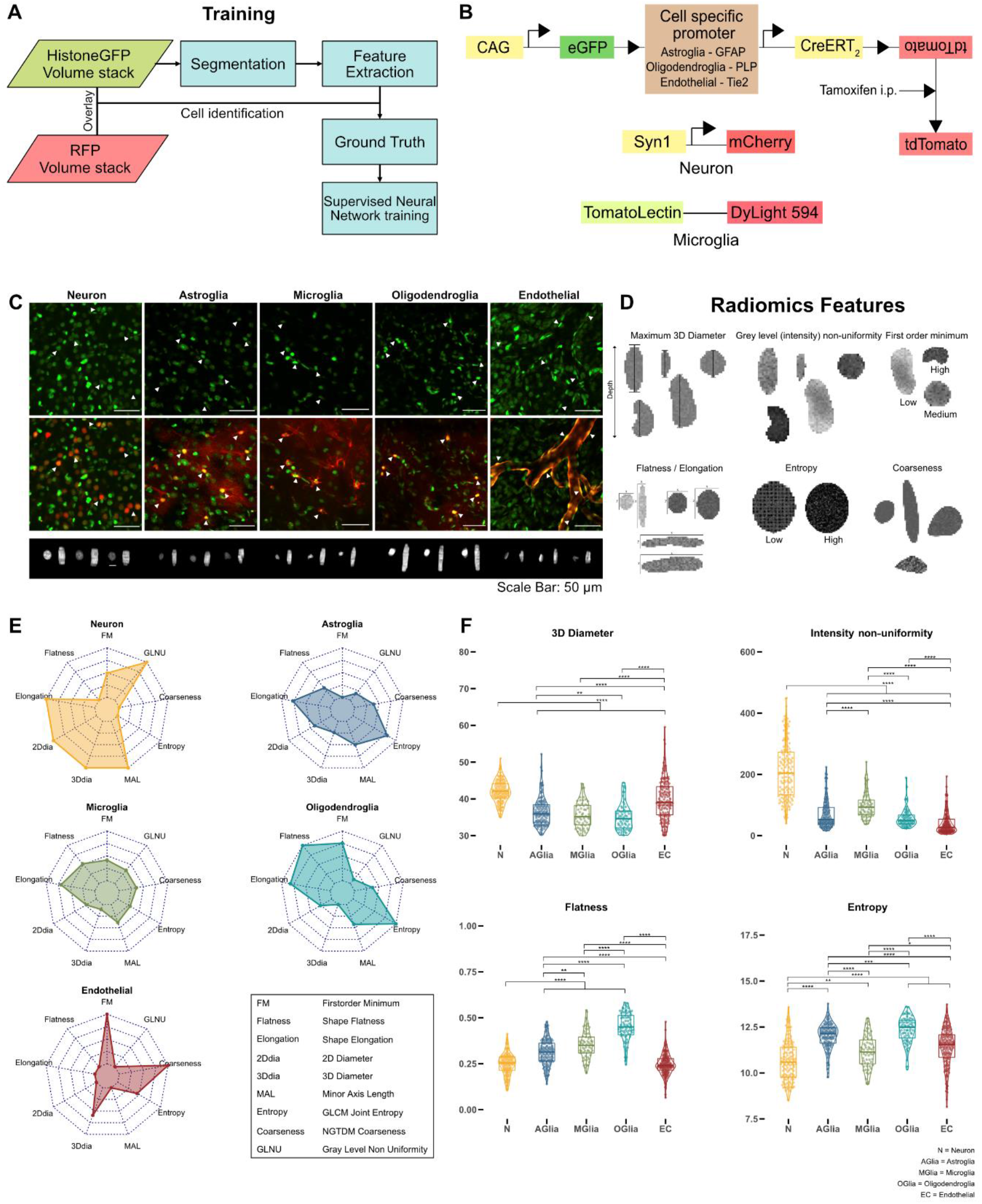
Strategies for training a neural network classifier using nucleus features. **(A)** Flow diagram showing classification training pipeline. Cell type specific red fluorescent proteins were used to manually identify H2B-eGFP nuclei. This information was used to train a classifier for each cell type. **(B)** Labelling strategies used for each cell type. Reporter mouse lines were created by breeding H2B-eGFP mice to carry a floxed sequence of the red fluorescence protein tdTomato and a tamoxifen inducible Cre-recombinase under the expression of different cell type specific promoters which were used for identification of astroglia, oligodendroglia and endothelial cells (see methods). Microglia and neurons were visualized using intracortical injections of Lycopersicon Esculentum (Tomato) lectin and an AAV expressing mCherry under the synapsin promoter respectively. **C)** Nuclei belonging to a specific cell type could be identified by a red fluorescent marker (maximum intensity projections; z = 20 µm). 3D-renderings of individual nuclei are shown in the lower panel. Scale bar: 50µm. **(D)** Illustrations of a subset of radiomics features showing examples of different shape, intensity and texture features. **(E)** Radar plots showing a subset of nuclear radiomics features for each cell type. **(F)** Comparison of nuclear features (3D diameter in voxels, (voxel resolution in xyz: 0.29 µm x 0.29 µm x 2µm), intensity non uniformity, flatness, entropy) between cell types (Wilcoxon test, Bonferroni correction for multiple comparisons, n of neurons: 135, n of astroglia: 137, n of microglia: 62, n of oligodendroglia: 72, n of endothelial cells: 155, p < 0.05 *, p < 0.01 **, p < 0.001 ***), N = Neuron, AGlia = Astroglia, MGlia = Microglia, OGlia = Oligodendroglia, EC = endothelial cells.

Automated segmentation of nuclei in the raw data was achieved using the StarDist neural network (Schmidt, Weigert et al. 2018), trained on manually traced and labelled ground truth datasets of the H2B-eGFP mouse line (Supplementary Figure 1H(i-ii),I). The trained network showed a high nucleus detection accuracy of 94 % as well as a good shape segmentation of nuclei (Supplementary Figure 1H (iii-iv), I). From the binary mask of individual nuclei and their respective pixel intensities in the raw image, in total 107 features were extracted using the PyRadiomics package (van Griethuysen, Fedorov et al. 2017), including 3D-diameter, flatness, grey level non-uniformity, entropy, first order minimum or coarseness (Figure 2D) (for a full list and short description of the features see Supplementary Tables 1 and 2). To reduce possible overfitting of the classification algorithm, a subset of 12 features were automatically selected using a sequential feature selection algorithm (Supplementary Table 2). Radar plots with these nuclear features visualize the differences between the cell types, for example neuronal nuclei having a larger 3D diameter and minor axis length in comparison to microglial and astroglial nuclei that exhibit a larger pixel entropy (Figure 2E). For certain features, significant differences between the cell types can be shown as well, for example flatness being significantly higher in nuclei of microglia than in nuclei of neurons (Figure 2F). Furthermore, combinations of two or three distinctive features allow for the visual separation of nuclei of different cell types in the 2D and 3D space (Supplementary Figure 3). When analyzing inhibitory and excitatory neurons, nuclei showed significant differences in shape and texture features (Supplementary Figure 2C (i, ii, iii, iv)).

Having demonstrated the ability to distinguish between cell types using only nuclear features, we created a neural network model for classifying cell types based on their features (Supplementary Figure 4A). Utilizing the dataset obtained from the five reporter mouse models, a single neural network classifier was trained for each cell type. The purpose of each classifier was to distinctly differentiate its corresponding cell type from the diverse array of other cell types (Figure 1C(i), 3A). To increase the amount of training data and equalize the nuclei counts for each cell type, thus reducing training bias, synthetic data was generated from the features of the original dataset (Supplementary Figure 4C). Synthetic data distribution fit well to the distribution of the original data for each cell type (Supplementary Figure 4C, D). After each nucleus was classified by all trained classifiers, it was either assigned to a single class (neurons, astrocytes, microglia, oligodendroglia, endothelial cells), two or more classes (undecided) or to none (unclassified). Precision and recall rates for the model were high for neurons and endothelial cells (Figure 3B). Due to their relative similarity, glial cells exhibited lower precision and recall rates. The classification accuracy for the entire training dataset was highest for neurons (98 %) and endothelial cells (99 %), whereas the classifier showed slightly lower accuracies for all glial cell types, especially for microglia (96 %) (Figure 3C, D, for the respective confusion plots see Supplementary Figure 5). To be able to distinguish between inhibitory and excitatory neurons, another classifier was trained on the ground truth data from excitatory and inhibitory cells. The classifier had a 93% overall accuracy (Supplementary Figure 2D), enabling a good distinction between inhibitory and excitatory neuronal cells. The 3D classification results for one example cortical volume containing 20123 nuclei (Supplementary Figure 6A, B) visualizes the relative abundance and positioning of cell types within the volume and their mutual spatial relationship (see also Supplementary Movie 1 for an animated version of the segmentation and classification process).

**Figure 3.**
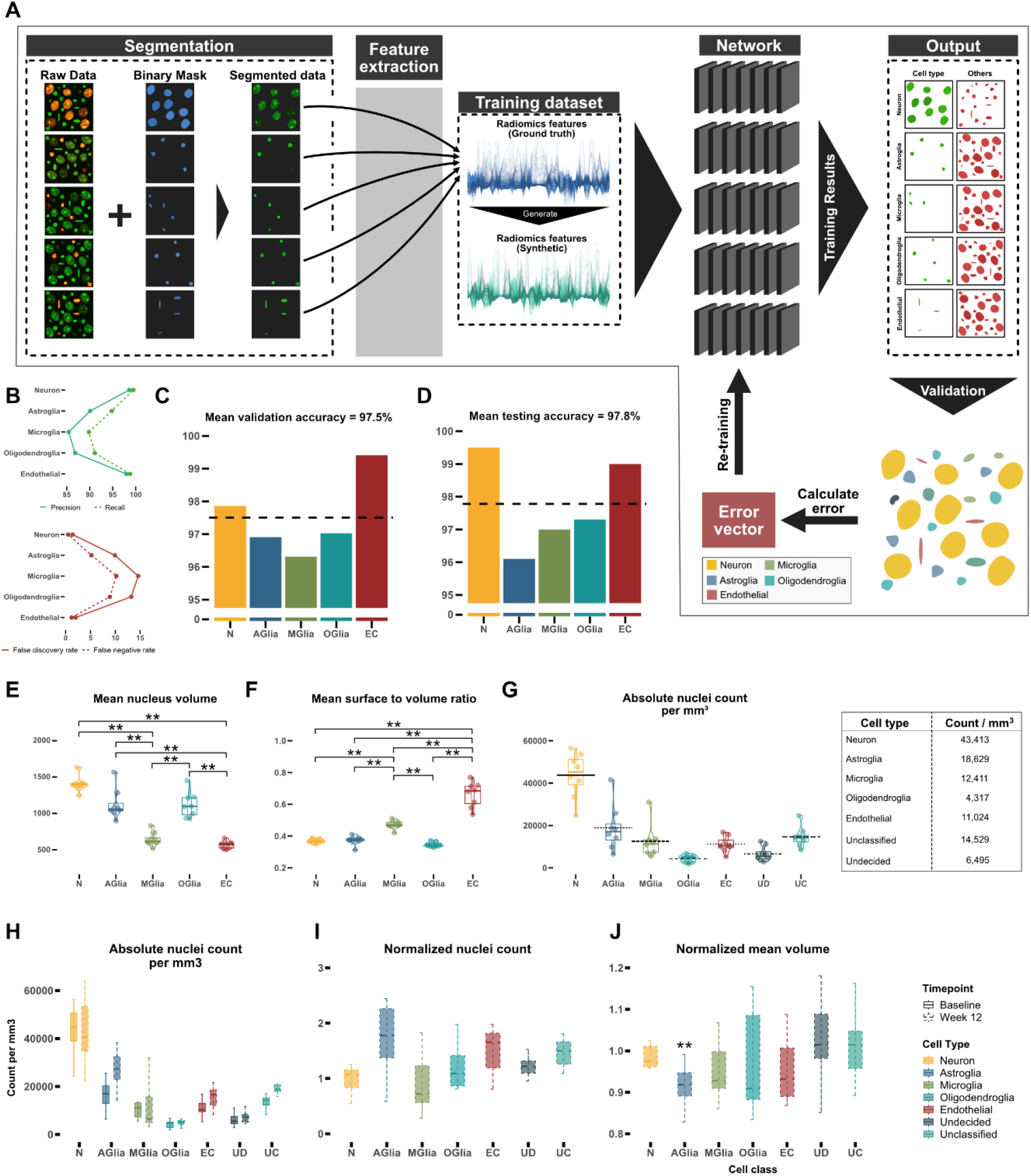
Training a neural network for cell type classification. **(A)** Schematic depiction of the training process. After segmenting nuclei of the raw data with StarDist for every cell type, radiomics features of nuclei were extracted from the ground truth data (blue) and synthetic data (green) was generated. For each cell type, a single neuronal network was trained to distinguish the corresponding nucleus type from the other nuclei. **(B)** Precision, recall, false discovery rate and false negative rates for the test dataset. **(C, D)** Mean accuracy for all cell types in the validation dataset (15 % of the ground truth dataset) and the test dataset (15 % of the ground truth dataset). **(E, F)** Comparison of mean nucleus volume (unit: µm^3^) and mean nuclear surface to volume ratio (unit: µm^-1^) of each cell type for classified volumes. **(G)** Number of nuclei per mm³ in the secondary motor cortex. **(H)** Number of nuclei per mm³ in the secondary motor cortex 12 weeks apart. **(I)** Comparison of normalized number of nuclei after 12 weeks. **(J)** Mean nucleus volume after 12 weeks normalized to baseline. (Significance testing for C, D, E, F, G, H, I, J: Wilcoxon test, p-values were corrected for multiple comparisons using the Bonferroni-method, n = 8 mice, p < 0.05*, p < 0.01 **, p < 0.001 ***), N = Neuron, AGlia = Astroglia, MGlia = Microglia, OGlia = Oligodendroglia, EC = endothelial cells.

To apply NuClear in a longitudinal way in mice, we studied 16 imaging volumes from 8 mice (male, 12-14 weeks at baseline) in layer 2/3 of the secondary motor cortex 12 weeks apart. When the trained model was applied, two features not chosen by the automated feature selection algorithm differed significantly between the classes: neuronal nucleus volume (Figure 3E) and endothelial mean surface to volume ratio, a measure for the sphericity of the nucleus (Figure 3F). Both findings are expected for each of the cell types and hence demonstrate that even features that were not used for training the neural network model are able to differentiate between the classes. At baseline a density of 43000 neurons per mm³ and of 57000 glial cells per mm³ (including unclassified cells, that were counted as glial cells) could be calculated, resulting in a glia to neuron ratio of 1.3 (Figure 3G). The density of excitatory neurons was 37000 cells/mm³, compared to 6000 cells/mm³ for inhibitory neurons (Supplementary Figure 2E). The density of cells 12 weeks after baseline imaging remained unchanged after correcting for multiple comparisons (Figure 3H). When normalizing these counts to baseline, only the number of astrocytes shows a trend to increase (Figure 3I), which could be either due to astrogliosis caused by aging or by continued reaction to the chronic window implantation. Interestingly, astroglial nuclear volume significantly decreased over time (Figure 3J), which might be due to altered transcriptional activity known to occur in astrocytes with aging (Boisvert, Erikson et al. 2018, Clarke, Liddelow et al. 2018), whereas the nuclei volume in inhibitory and excitatory neurons did not change over time (Supplementary Figure 2F). This latter result illustrates that our method can extract even subtle and unexpected changes from of a vast parameter space of different aspects of tissue composition.

## Discussion

Here, we present a novel method for intravital, longitudinal, comprehensive, large scale and 3D determination of tissue composition based on a deep learning algorithm trained on radiomics features of cell nuclei (NuCLear -- Nucleus-instructed tissue composition using deep learning). We demonstrate that cell types and even subtypes can be reliably distinguished via the respective properties of their cell nucleus and that an inventory of cells and their 3D position can be generated for a large imaging volume. To demonstrate the usability of NuCLear, we analyzed volumetric images of the cortex of H2B-eGFP mice acquired with in vivo two photon microscopy. To be able to image a large dataset in a comparatively short time, we chose an imaging resolution with a z-step size of 2 µm and a lateral resolution of 0.3 µm. We were able to image a whole 3D volume of 700 µm³ consisting of more than 20000 cells in around 20 minutes, making it possible to perform large scale data acquisition in a repeated manner, enabling longitudinal intravital monitoring of tissue histology over time and in response to perturbations, treatments, or adaptive processes such as learning or exploring.

Neurons and endothelial cells showed the best classification results regarding total accuracy and precision as well as recall rates (Figure 3B), as their nucleus feature profile turned out to be markedly different between each other as well as between the individual glial cell types (Figure 2E, F). Differentiation between the three different glial cell types was more difficult to achieve, resulting in lower overall accuracy rates as well as lower precision and recall rates, due to the higher similarity of their respective nucleus feature profiles. Nevertheless, with an accuracy above 90% our approach can reliably be used to subclassify glia. We expect that a higher imaging resolution as well as more training data would further increase the classifier performance. Another reason for the comparatively lower classification accuracy in microglia might be due to acute injection of tomato-lectin to label these cells, which could have introduced a bias towards activated microglia. Nevertheless, the overall accuracy of the microglia classifier for the whole training dataset was around 96%, owing to the fact, that non-microglial nuclei could be identified correctly. For astrocytes, we used GFAP-labelled cells for classification, a label that primarily marks reactive astrocytes and has a higher expression in older animals. Future studies could optimize this labelling strategy. Additional confirmation of the classifier’s precision can be inferred by examining characteristics of the nuclei, like average nuclear volume and the surface-to-volume ratio, which were not utilized in the training phase (Figure 3E, F). These features showed distinct clustering and low standard deviation in each class as well as significant differences between the cell types.

When applying the NuCLear to imaging volumes from Layer 2/3 of the secondary motor area and calculating the number of cells per mm³, a marked similarity of the number of neurons with published results is evident (43413 ± 1003 vs 45374±4894) (Ero, Gewaltig et al. 2018) (Figure 3G). Astrocyte numbers are similar (18629 ± 1007 vs 13258± 1416), microglia numbers differ more (12411 ± 745 vs 24584.6± 2687). Oligodendrocyte numbers seem to be vastly different (4317 ± 154 vs 51436.1± 488), which might stem from the fact that only Layer 2/3 was analyzed, excluding deeper cortical layers that have been shown to contain most oligodendrocytes (Tan, Kalloniatis et al. 2009). When all glial cells as well as undecided and unclassified cells are summed up, a Glia/Neuron ratio of 1.3 can be calculated, a result that is in line with known results of the rodent cortex (Herculano-Houzel 2014). Candidates for unclassified cells could be the relatively large population of oligodendrocyte precursor cells (about 5% of all cells in the brain (Dawson, Polito et al. 2003)). After classifying the neurons into subtypes, 86% of all neurons were classified as excitatory and 14% as inhibitory (37199±1387 vs 6214±1001 respectively), which is in par with previous studies that reported 10-20% of neurons in layer 2/3 of the cortex being inhibitory interneurons (Kummer, Mitric et al. 2020, Algamal, Russ et al. 2022). In conclusion, cell type counts are comparable to published results, supporting the validity of our approach.

A decisive advantage of NuCLear over existing ex-vivo methods is the ability to study cell type changes over time in the same imaging volume and thus achieving a higher statistical power in experiments with fewer animals, which is crucial for complying with the “3R” rules in animal research (reduce, replace, refine). We succeeded to image selected locations up to 1 year after the implantation of chronic cranial windows. When analyzing the same imaging volumes from the secondary motor cortex 12 weeks apart, we were able to detect a trend towards astrogliosis, which might be due to the aging process leading to a higher GFAP-reactivity (Palmer and Ousman 2018). The significant decrease in mean nucleus volume of astroglia over time could be attributed to altered transcriptional activity associated with aging (Boisvert, Erikson et al. 2018, Clarke, Liddelow et al. 2018) since astroglial reactivity changes in the cortex over time (Clarke, Liddelow et al. 2018). Another factor underlying astrogliosis might arise from the continued presence of the chronic cranial window (Guo, Zou et al. 2017). Our study illustrates only one possible application of NuCLear and subsequent analysis of 3D tissue composition. Here we focused on the identification of cell types, but the 3D datasets generated allow for additional analyses such as statistical distribution of cell types relative to each other, their nearest neighbor distances or inferences on physical tissue volume (Asan, Falfan-Melgoza et al. 2021). Our classifiers mostly rely on geometrical parameters such as nucleus shape, but also consider texture information to a certain degree. Pathophysiological conditions that alter nuclear shape and texture might influence classification accuracy. An increased sampling resolution may further increase textural information and thereby yield even higher classification accuracies. For now, segmentation and classification depend on the type of microscope used, quality of the images, depth of imaging as well the region which was analyzed. To make the classification more robust and accountable for different qualities of fluorescence signal, augmentations to the training data could be added such as 3D blurring with PSF-shaped convolutions.

NuCLear will be applicable to different organs as techniques have emerged in the last years to perform in vivo imaging through chronic windows in other rodent organs such as skin, abdominal organs, the tongue or spinal cord (Choi, Kwok et al. 2015), depending on the image quality and the possibility to acquire images during multiple timepoints. Other organisms and in vitro preparations like cell cultures or organoids can also be utilized, provided that the respective ground truths can be generated. It will be usable with a variety of microscopy techniques such as confocal-, light-sheet- or three-photon microscopy, making large-scale tissue analysis much more accessible. We assume that it will be possible to detect more cell types by adding classifiers trained with appropriate ground truth data. In this way, we propose a readily usable method to implement large-scale tissue analysis ex-vivo or in-vivo to study effects of interventions on cell type compositions of different organs.

## Materials and Methods

### Ethical approval

The entire study was conducted in line with the European Communities Council Directive (86/609/ECC) to reduce animal pain and/or discomfort. All experiments followed the German animal welfare guidelines specified in the TierSchG. The study has been approved by the local animal care and use council (Regierungspräsidium Karlsruhe, Baden-Wuerttemberg) under the reference numbers G294/15 and G27/20. All experiments followed the 3R principle and complied with the ARRIVE guidelines (Percie du Sert, Hurst et al. 2020).

### Animals

Adult transgenic mice expressing the human histone 2B protein (HIST1H2B) fused with enhanced green fluorescence protein (eGFP) under the control of the chicken beta actin promoter (CAG-H2B-eGFP) (Hadjantonakis and Papaioannou 2004) (B6.Cg-Tg(HIST1H2BB/EGFP)1Pa/J, Jackson Laboratory; # 006069) were used in all animal experiments. For the reporter mouse lines, H2B-eGFP mice were bred to carry a floxed sequence of the red fluorescent protein tdTomato and a tamoxifen inducible Cre-recombinase under the expression of different cell type specific promoters: GFAP (Glial Fibrillary Acidic Protein) for astroglia (HIST1H2BB/EGFP-GFAP/ERT2CRE-CAG/loxP/STOP/loxP/tdTomato), PLP (Proteolipid Protein) for oligodendroglia (HIST1H2BB/EGFP-PLP/ERT2CRE-CAG/loxP/STOP/loxP/tdTomato) and Tie2 (Tyrosine Kinase) for endothelial cells (HIST1H2BB/EGFP-Tie2/ERT2CRE-CAG/loxP/STOP/loxP/tdTomato). Expression of the Cre-recombinase was achieved over a course of up to 5 days with daily intraperitoneal injection of two doses of 1mg Tamoxifen (Sigma Life sciences; dissolved in 1 part ethanol (99.8% absolute) and 10 parts sunflower oil) (Figure 2B, Supplementary Figure 1D). Microglia were visualized using intracortical injection of Lycopersicon Esculentum (Tomato) lectin coupled with DyLight 594 during the cranial window surgery after dilution by 1:39 in 150mM NaCl, 2.5 mM KCl, 10mM HEPES at pH 7.4 (Schwendele, Brawek et al. 2012) (Figure 2B, Supplementary Figure 1D). For neuronal labelling, an in house produced adeno-associated virus expressing mCherry under the Synapsin promotor was cortically injected during the cranial window surgery (Figure 2B, Supplementary Figure 1D). To label excitatory and inhibitory neurons, a viral labelling strategy was implemented via intracortical injections, using AAV5-CamKIIα-mCherry (Addgene plasmid 114469) and AAV1-mDLX-mRuby2 (Addgene plasmid 99130) (Supplementary Figure 2) (Chan, Jang et al. 2017). All mice were between 10-18 weeks old during baseline imaging (5 female/10 male).

### Chronic cranial window implantation

A craniectomy procedure was performed on each mouse to enable in vivo two-photon imaging following a predefined protocol as described before (Holtmaat, Bonhoeffer et al. 2009) (Supplementary Figure 1A). Briefly, the mice were anesthetized with an intraperitoneal injection (i.p.) of a combination of 60 µl Medetomidine (1mg/ml), 160 µl Midazolam (5 mg/ml) and 40 µl Fentanyl (0.05 mg/ml) at a dosage of 3 µl/g body weight. The head was shaved, and the mice were fixed with ear bars in the stereotactic apparatus and eye ointment was applied. Xylocain® 1 % (100 µl, Lidocaine hydrochloride) was applied under the cranial skin and 250 µl Carprofen (0.5 mg/ml) was injected subcutaneously (s.c.). The skin was removed to expose the skull and the surface of the skull was made rough to allow the cement to adhere better. A skull island (approx. 6mm Ø) was drilled centered at 1 mm rostral to bregma using a dental drill and removed with a forceps (#2 FST by Dumont) making sure not to damage the cortical surface. For improved imaging condition, the dura was carefully removed from both hemispheres using a fine forceps (#5 FST by Dumont). Normal rat ringer solution (NRR) was applied on the exposed brain surface to keep it moist, and a curved cover glass was placed on top to cover it (Asan, Falfan-Melgoza et al. 2021). With dental acrylic cement (powder: Paladur, Kulzer; activator: Cyano FAST, Hager & Werken GmbH) the cover glass was sealed, and excess cement was used to cover the exposed skull and edges of the skin. A custom designed 3-D printed holder was placed on top of the window and any gaps were filled with cement to ensure maximum adhesion with the skull. After the procedure, the mice were injected (i.p. / s.c.) with a mix of 30 µl Atipamezole (5 mg/ml), 30 µl Flumazenil (0.1 mg/m) and 180 µl Naloxon (0.4 mg/ml) at a dosage of 6 µl per gram body weight. To ensure proper recovery of the mice, 3 more doses of Carprofen were given every 8-12 hours and the mice were placed onto a heating plate and monitored.

### In vivo two photon imaging

Imaging was carried out with a two-photon microscope (TriM Scope II, LaVision BioTec GmbH) with a pulsed Titanium-Sapphire (Ti:Sa) laser (Chameleon Ultra 2; Coherent) at excitation wavelengths of 860nm for DyLight 594 and 960nm for H2GFP, tdTomato and mCherry respectively. A water immersion objective (16x; NA 0.8, Nikon) was used to obtain volumetric 3D stacks. Individual frames consisted of 694 µm x 694 µm in XY with a resolution of 0.29 µm/pixel. Stacks were obtained at varying depths up to 702 µm from the cortical surface with a step size of 2 µm in Z. Prior to each imaging session, the laser power was adjusted to achieve the best signal to noise ratio. Adaptations were made to minimize the effect of laser attenuation due to tissue depth by creating a z-profile and setting the laser power for different imaging depths while making sure to minimize oversaturation of pixels.

Mice were initially anesthetized using a vaporizer with 6% isoflurane, eye ointment was applied, and mice fixed in a custom-built holder on the microscope stage. Isoflurane was adjusted between 0.5 – 1.5% depending on the breathing rate of each mouse to achieve a stable breathing rate of 55 – 65 breaths per minute with an oxygen flow rate of 0.9 – 1.1 l/min. A heating pad was placed underneath the mouse to regulate body temperature. An infrared camera was used for monitoring of the mice during the imaging.

For ground truth training data reporter mouse lines for neurons and neuronal subtypes, astroglia, oligodendroglia and endothelial cells were imaged 2-4 weeks after the cranial window surgery. Microglia reporter mice were imaged immediately after the cranial window surgery for a minimal inflammatory reaction (for a full list of reporter mice numbers see Supplementary Figure 4G). For the data obtained from adult H2B-eGFP mice used for classification, 3D volumetric stacks were imaged in the secondary motor cortex at two different timepoints. This included baseline imaging which was performed at 3-4 weeks after the chronic cranial window surgery and 12 weeks after the baseline timepoint. To investigate a possible age effect, mice underwent a sham surgery at the left hind paw 1 week after baseline surgery as part of a different study.

### Automatic nuclei segmentation using StarDist

Automated nuclei segmentation was achieved using the StarDist algorithm, which implements a neural network to detect star-convex polygons in 2D and 3D fluorescence imaging (Schmidt, Weigert et al. 2018). The StarDist model was trained to segment nuclei in 3D using an in-house developed Jupyter Notebook. A GUI based version of the software for training of the segmentation (NucleusAI) can be found here: https://github.com/adgpta/NuCLear. Nuclei were manually traced and segmented using the segmentation editor in Fiji (Schindelin, Arganda-Carreras et al. 2012) (Supplementary movie 2). In total, 15 different crops from the two-photon volumetric data with approximately 150 – 200 nuclei each were used for training the StarDist 3D segmentation classifier (Supplementary Figure 1I).

Detection accuracy and precision were calculated after visualization of raw data and segmentation in ImageJ/Fiji. True positive nuclei (TP), false positive nuclei (FP) and false negative nuclei (FN) were manually counted. 4 arbitrary volumetric images were selected containing up to 80 nuclei per crop. Accuracy was calculated as TP / (TP + FP + FN). Precision was calculated as TP / (TP + FP).

### Feature extraction of segmented nuclei using PyRadiomics

Using the PyRadiomics python package (van Griethuysen, Fedorov et al. 2017), in total 107 radiomics features were extracted for each nucleus after segmentation using StarDist, including, but not limited, to morphological features, intensity features and texture features (see Supplementary table 1). Nuclei touching the borders of the imaging volume were excluded.

### Extraction of data for training of the classifier

To perform supervised training of the deep neural network for cell type classification, a ground truth dataset was created using the two-photon volumetric fluorescence data, automatically segmented nuclei, which were assigned a unique label, and the radiomics features. Using the red fluorescence channel, nuclei belonging to a specific cell type were manually identified after creating a composite image of the green and red fluorescence images (Supplementary Figure 4B). This allowed for identification of the label by synchronizing the composite image and the StarDist output in ImageJ/Fiji and extraction of radiomics features for these individual nuclei. Approximately 70-400 nuclei were identified for each cell type (Supplementary figure 4G).

To increase the number of nuclei for training and make it equal in size for all cell types and thus avoid a bias in training, synthetic datasets were created from nuclei features using the synthetic data vault (SDV) package (Patki, Wedge et al. 2016), which utilizes correlations between features of the original dataset as well as mean, minimum or maximum values and standard deviations. The synthetic data generated by the model fit to the original dataset with approximately 83% accuracy (Supplementary Figure 4D).

### Training of the classification model

All training and validation of the classifiers were performed with a custom MATLAB (R2021a, Mathworks) script, visualization was performed in ImageJ/Fiji (Supplementary movie 3). Original and synthesized data were merged into a single table and random indices were generated to divide the dataset into a training, test, and validation dataset with 70%, 15% and 15% of the total dataset, respectively. Datasets with different amounts of synthetic data combined with the original dataset were created and training accuracy was later compared between them: orig = original dataset, orig * 2.5 = dataset containing 2.5 times the amount of data as the original dataset, orig * 2.5 (down sampled) = dataset down sampled to minimum sample count (after 2.5 fold increase) to equalize sample numbers for all cell types, orig * 9 = dataset containing 9 times the amount of data as the original dataset, orig * 9 (down sampled) = dataset down sampled to minimum sample count (after 9 fold increase) to equalize sample numbers for all cell types (Supplementary Figure 4E, F). Down sampled datasets were created by selecting the class with the minimum number of nuclei and removing random samples from the other classes to match the nuclei count of the selected class. The same training, test, and validation datasets were used for training five different classifiers, one for each cell type. In the training dataset, for each classifier, the class of interest was assigned a unique identifier and all other cell classes were denoted by another identifier, for example, when training the neuronal classifier, the neurons were labelled as “Neuron” while all the other cell types in the dataset were labelled “Other” (Figure 1C(i), Figure 3A, Supplementary Figure 4A). Each classifier had the same design. Initially, a sequential feature selection algorithm “sequentialfs” was applied to the whole dataset to extract 12 features with the highest variability to reduce data overfitting. These were fed into the “feature input layer” of the classifier. The features were z-score normalized for better compatibility between each class. A “fullyconnected” layer of 50 outputs was created and a mini-batch size of 16 samples was set to reduce the time and memory consumption during training. A batch normalization layer was added to reduce sensitivity to the initialization of the network. The activation layer ReLU (Rectified Linear Unit) was added for thresholding operations. Another “fullyconnected” layer was added, which sets the number of outputs identical to the number of input classes and a second activation layer “softmax” was chosen for classifying the output into the two separate groups. The “adam” adaptive moment estimation optimizer (Kingma and Ba 2017) was selected as the solver for the training network which updates the network weights based on the training data in an iterative manner. The training data and the validation data were shuffled every epoch, i.e. a complete pass or iteration through the entire training dataset during the training process, before training and network validation, respectively. After each classifier was trained, further automatic and visual validation was performed to check for accuracy.

### Classification of the H2B-eGFP data

After acquiring volumetric images of H2B-eGFP nuclei (n=8 mice, male), automated segmentation of nuclei was performed with the StarDist neuronal network (see above, Supplementary Figure 6A, C). Features for the segmented nuclei were extracted using PyRadiomics (see above) and then classified with all the different classifiers trained on the orig * 9 dataset (see “Training of the classification model” for a description of the dataset). Thus, a single nucleus would be either labelled as belonging to one of the five classes or to the “Other” class. Nuclei that were assigned to multiple classes, were labelled as “Undecided” (UD). Any nuclei that were identified as “Other” by all the classifiers were labelled as “Unclassified” (UC). Nuclei of neurons were further subdivided into excitatory and inhibitory subtypes.

### Statistical analysis

All classified nuclei and their features were stored in a local MySQL database (MySQL Workbench 8.0). The data were imported into R (R. Core Team 2021) for statistical analysis. Cell count, mean nuclei volume and first nearest neighbor distance were calculated for each cell type for different timepoints. Since the distribution of the data was non-parametric, the Wilcoxon signed-rank test was used for all statistical testing and p-values were corrected for multiple comparisons using the “bonferroni”-method. Plots were created using the R package “ggplot2” (Wickham 2016).

## Contributions

ADG: Data acquisition, implementation and development of segmentation and classification pipeline and scripts, training of classifiers, conceptual inputs, generation of figures, writing of the MS.

LA: Acquisition of experimental ground truth data with reporter mouse lines using 2PLSM imaging and establishment of tomato-lectin labelling, proof-of-concept with initial development of solutions for segmentation and classification.

CAB: Development of segmentation, application of StarDist, contributions to classification. JJ: Data acquisition, training of StarDist for segmentation.

TK: Initial project idea and concept, writing of the MS, supervision.

JK: Conceptualization, implementation, experimentation, analysis, writing of the MS, project management and supervision.

## Data and code availability

All datasets used in the study are available on the heiData archive: https://heidata.uni-heidelberg.de/dataset.xhtml?persistentId=doi:10.11588/data/L3PITA

Code and exemplary data can be found on the github page: https://github.com/adgpta/NuCLear

## Acknowledgements

We thank Michaela Kaiser for her excellent technical support during preparation and execution of experiments. We thank Sidney Cambridge for devising breeding schemes for mouse lines needed for ground truth generation and for help with mouse breeding. We gratefully acknowledge the support by the German Research Foundation (DFG) (SFB1158, project B08 awarded to TK), the data storage service SDS@hd, supported by the Ministry of Science, Research and the Arts Baden-Württemberg (MWK) and the DFG through grant INST 35/1314-1 FUGG, as well as the high-performance cluster bwForCluster MLS&WISO, supported by the MWK and the DFG through Grant INST 35/1134-1 FUGG.

---

**Supplementary figure 1.**
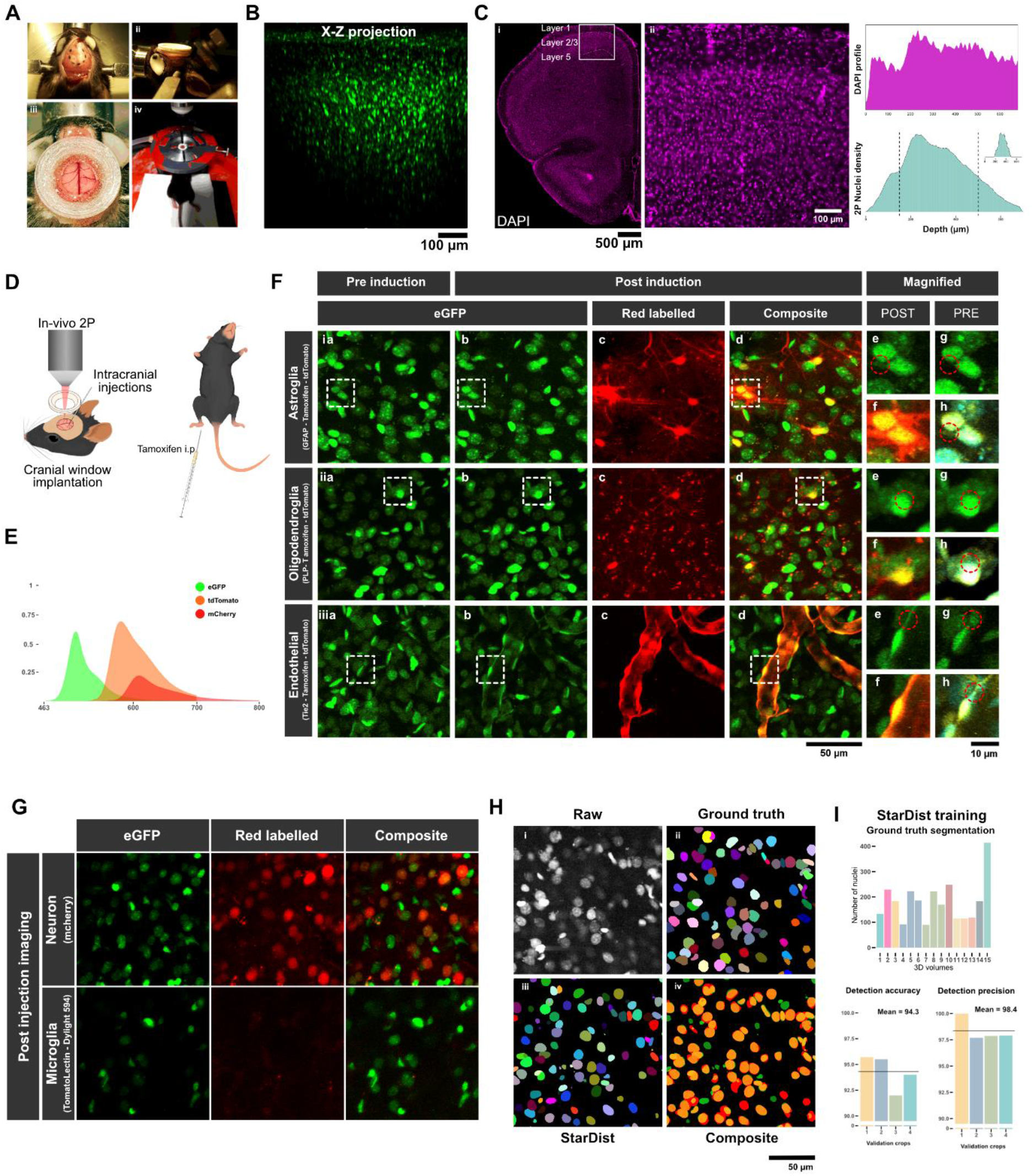
**(A) (i,ii,iii)** Chronic cranial window implantation using a curved glass cover slip and a custom 3D printed holder. **(iv)** Fixation of the anesthetized mouse in the custom holder for imaging under the 2-photon microscope. **(B)** X-Z maximum intensity projection of a two-photon volumetric stack (700µm x 700µm x 700µm) **(C)** Imaging position on a DAPI stained brain slice. Slice thickness 50 µm. DAPI intensity profile showing similar distribution of intensity as a two-photon image of nuclei. The decreasing nucleus density in the two-photon image stack is a result of attenuation of fluorescence signal at higher depths. Inlay: Nucleus density distribution of sub-volumes used for data analysis. **(D)** Scheme showing intracranial and intraperitoneal injection strategies for labelling cell types. **(E)** Fluorescence emission spectra for eGFP, tdTomato and mCherry (y axis: fluorophore emission normalized to quantum yield, source: fpbase.org) **(F)** Labelling strategies for red fluorescence expression in reporter mouse lines. Visualization of crosstalk between the eGFP and tdTomato signal after induction with tamoxifen. Overlay of pre (cyan) and post (yellow) GFP after image alignment with the ImageJ plugin bUnwarpJ (Arganda-Carreras, Sorzano et al. 2008). **(G)** No crosstalk is visible between GFP and mCherry signals (upper panel) or eGFP and Tomato lectin-Dylight 594 signals (lower panel). **(H)** (i) Raw data of H2B-eGFP signal (ii) manually labelled ground truth (iii) StarDist segmentation (iv) composite image of ground truth (red) and StarDist segmentation (green). **(I)** Upper panel: Count of manually segmented ground truth nuclei for StarDist training, each color depicts an individual mouse. Lower panel: StarDist nucleus detection accuracy and precision, each bar represents an individual imaging volume.

**Supplementary Figure 2.**
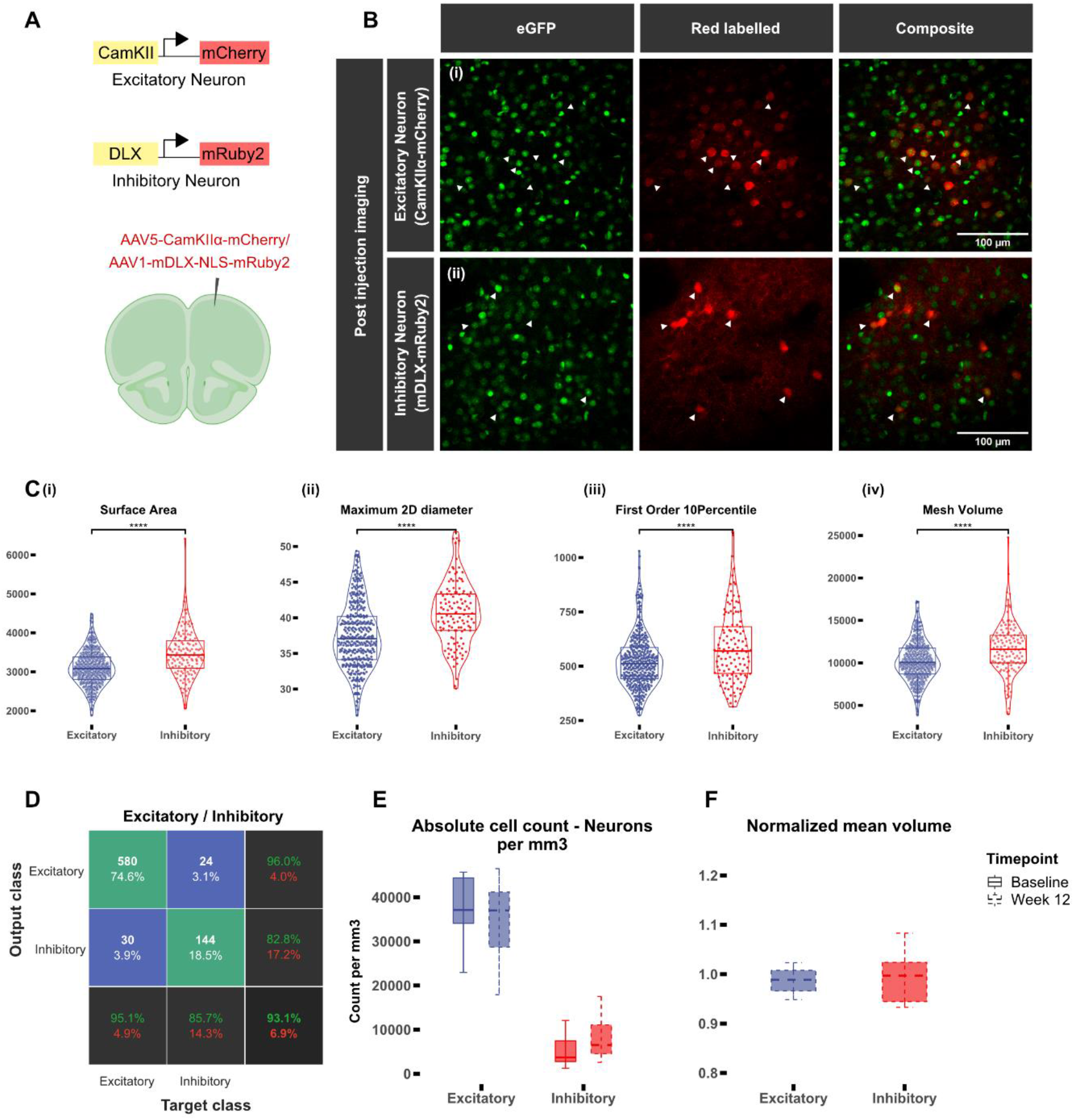
Training a classifier to distinguish excitatory from inhibitory neurons. **(A)** Labelling strategy for excitatory and inhibitory neurons. Intracortical injections were performed at ∼400µm deep from the cortex of H2B-eGFP mice using AAV5-CamKIIα-mCherry and AAV1-mDLX-NLS-mRuby2 to visualize excitatory and inhibitory neurons. **(B)** Post-injection images. Inhibitory neurons were imaged 2-3 weeks after injection, excitatory neurons were imaged 3-4 weeks after injection. **(C)** Radiomics features showing differences between excitatory and inhibitory neurons for (i) surface area (in pixel ² [resolution x = 0.29 µm, y = 0.29 µm, z = 2 µm]) (ii) maximum 2D diameter (in pixel, same resolution as in (i)) (iii) first order 10^th^ percentile (iv) mesh volume in voxels (resolution as in (i)), (n of excitatory neurons: 396, n = 3 mice, n of inhibitory neurons = 122, n = 2 mice). **(D)** Confusion plot for the classifier. Rows show the predicted class (output class), and the columns show the true class (target class). Green fields illustrate correct identification whereas blue fields illustrate erroneous identifications. The number of observations and the percentage of observations compared to the total number of observations are shown in each cell. Column on the far right shows the precision (or positive predictive value) and false discovery rate in green and red respectively. Bottom row denotes recall (or true positive rate) and false negative rate in green and red. Cell on the bottom right shows overall accuracy of the classifier. **(E)** Number of nuclei per mm³ in the secondary motor cortex at baseline after 12 weeks (dashed line) (n = 8 mice). **(F)** Mean nucleus volume after 12 weeks normalized to baseline (n = 8 mice). (Significance testing for C, E, F: Wilcoxon test, p-values were corrected for multiple comparisons using the Bonferroni-method, p < 0.05*, p < 0.01 **, p < 0.001 ***).

**Supplementary Figure 3.**
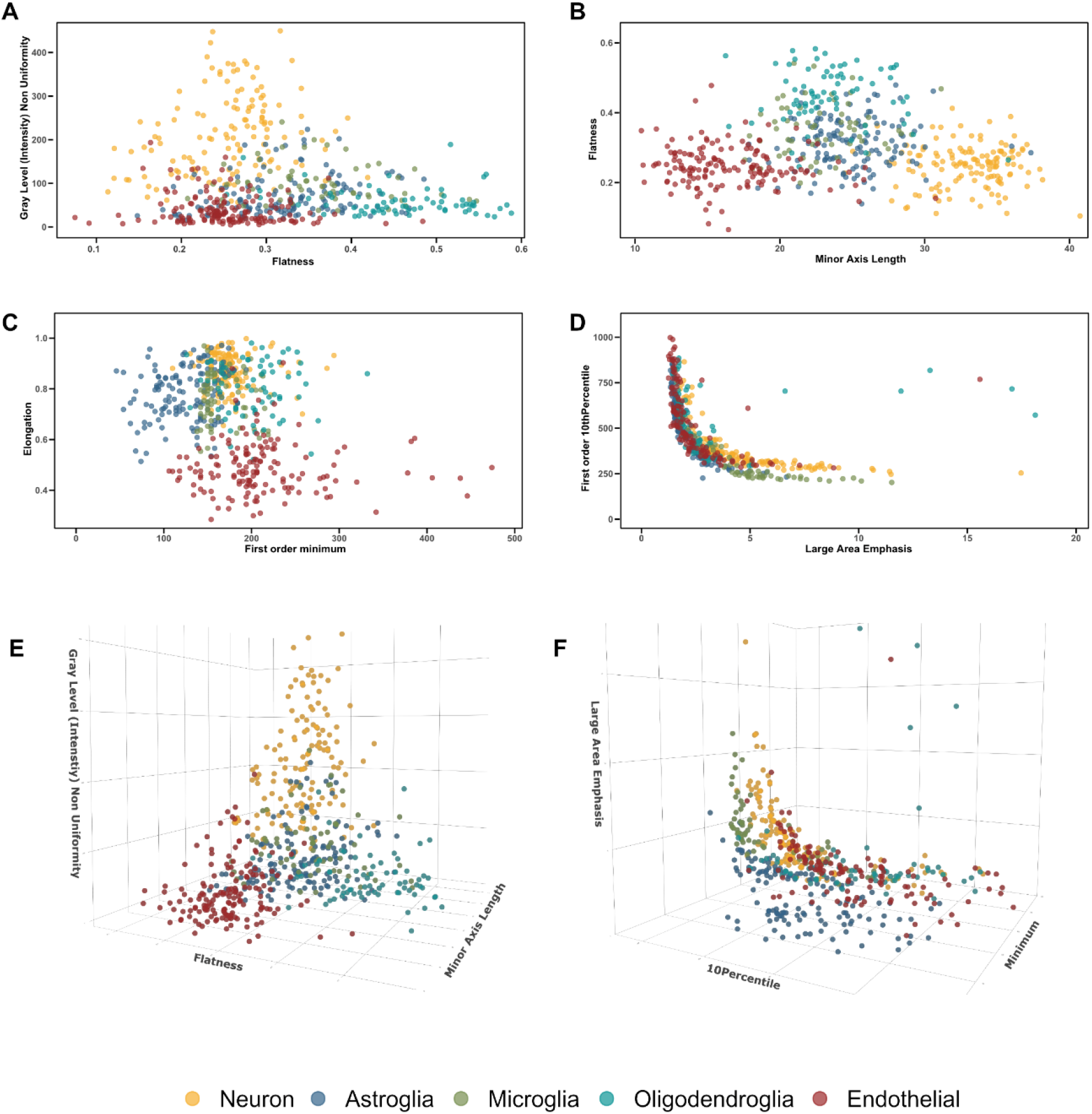
Combinations of nuclear morphology, intensity, and texture features show clear distinction between cell types. **(A)** Flatness, gray level non-uniformity. **(B)** Minor axis length, flatness. **(C)** First order minimum, elongation. **(D)** Large area emphasis, first order 10^th^ percentile. **(E, F)** Combinations of features in three dimensions.

**Supplementary Figure 4.**
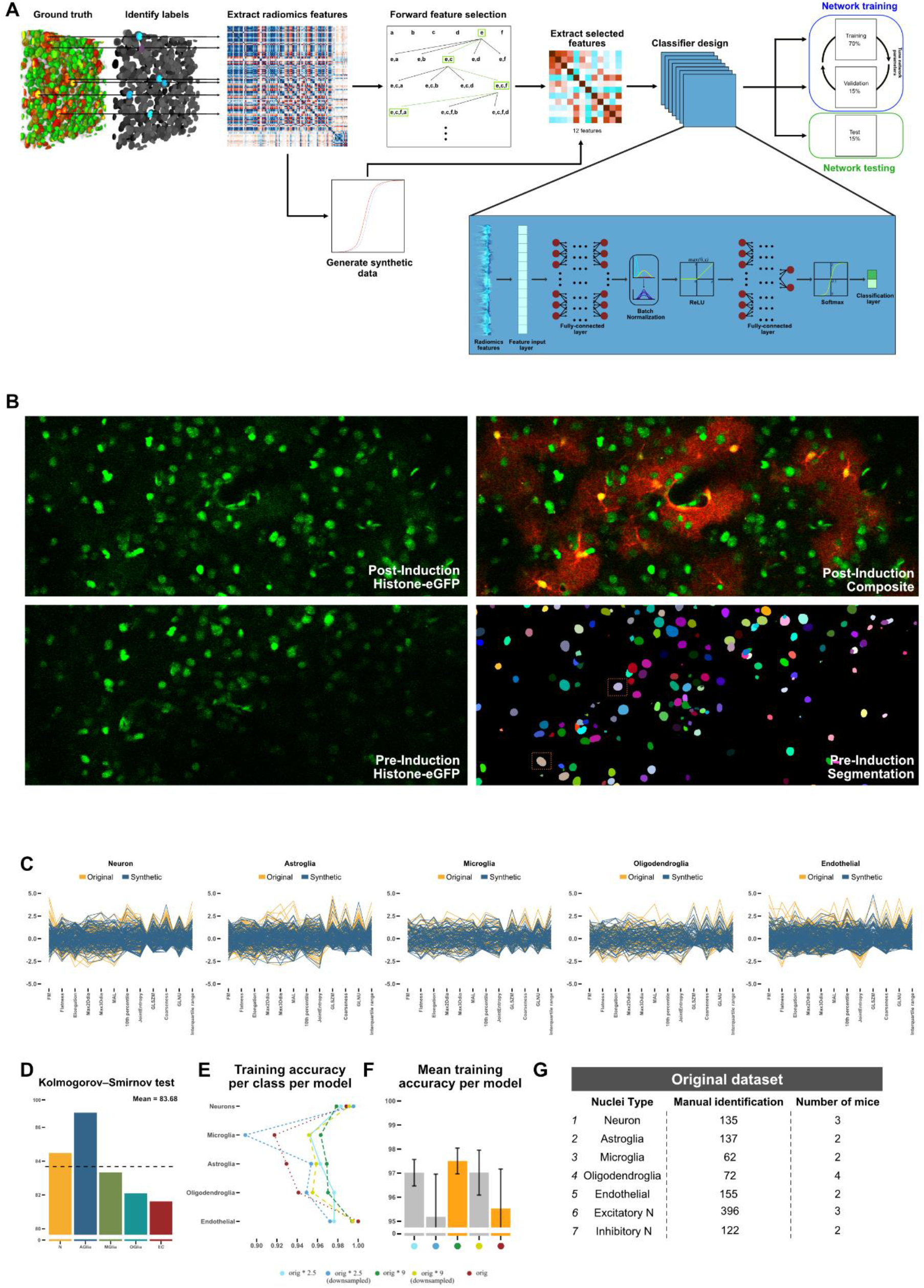
**(A)** Visualization of the entire classification training process. After ground truth data were selected, a sequential forward feature selection algorithm was applied to extracted features from all nuclei of all cell types, which selected 12 features from the 107 radiomics features. Synthetic data was generated from all the radiomics features of all nuclei and all cell types. The combined dataset was used to train a classifier with the training, validation and test data comprising 70%, 15%, and 15% of the combined dataset. **(B)** Example showing manual selection of ground truth data for supervised training. Post-induction green and red channels were overlayed to create a composite. Any nuclei appearing yellow (possessing green and red fluorescence) were identified in the pre-induction GFP channel and from its corresponding segmentation, a label id was acquired and later used to identify the extracted features. **(C)** Synthetic training data generated from the original dataset matched the features of the original datasets. **(D)** Statistical analysis of similarity between the distribution of original data and distribution of synthetic data (K-S test; mean = 83.68%). **(E)** Datasets with different amounts of synthetic data were created and training accuracy was compared between them, orig = original dataset, orig * 2.5 = dataset containing 2.5 times the amount of data as the original dataset, orig * 2.5 (down sampled) = dataset down sampled to minimum sample count (after 2.5 fold increase) to equalize sample numbers for all cell types, orig * 9 = dataset containing 9 times the amount of data as the original dataset, orig * 9 (down sampled) = dataset down sampled to minimum sample count (after 9 fold increase) to equalize sample numbers for all cell types. **(F)** Mean accuracy of classifiers trained 5 times with different combinations of synthetic data; error bars denote standard deviation (SD). light blue = orig * 2.5, dark blue = orig * 2.5 (downsampled), green = orig * 9, yellow = orig * 9 (down sampled), red = orig. **(G)** Table listing the amount of manually identified nuclei used in the training process (Excitatory N: excitatory neurons, Inhibitory N: inhibitory neurons).

**Supplementary Figure 5.**
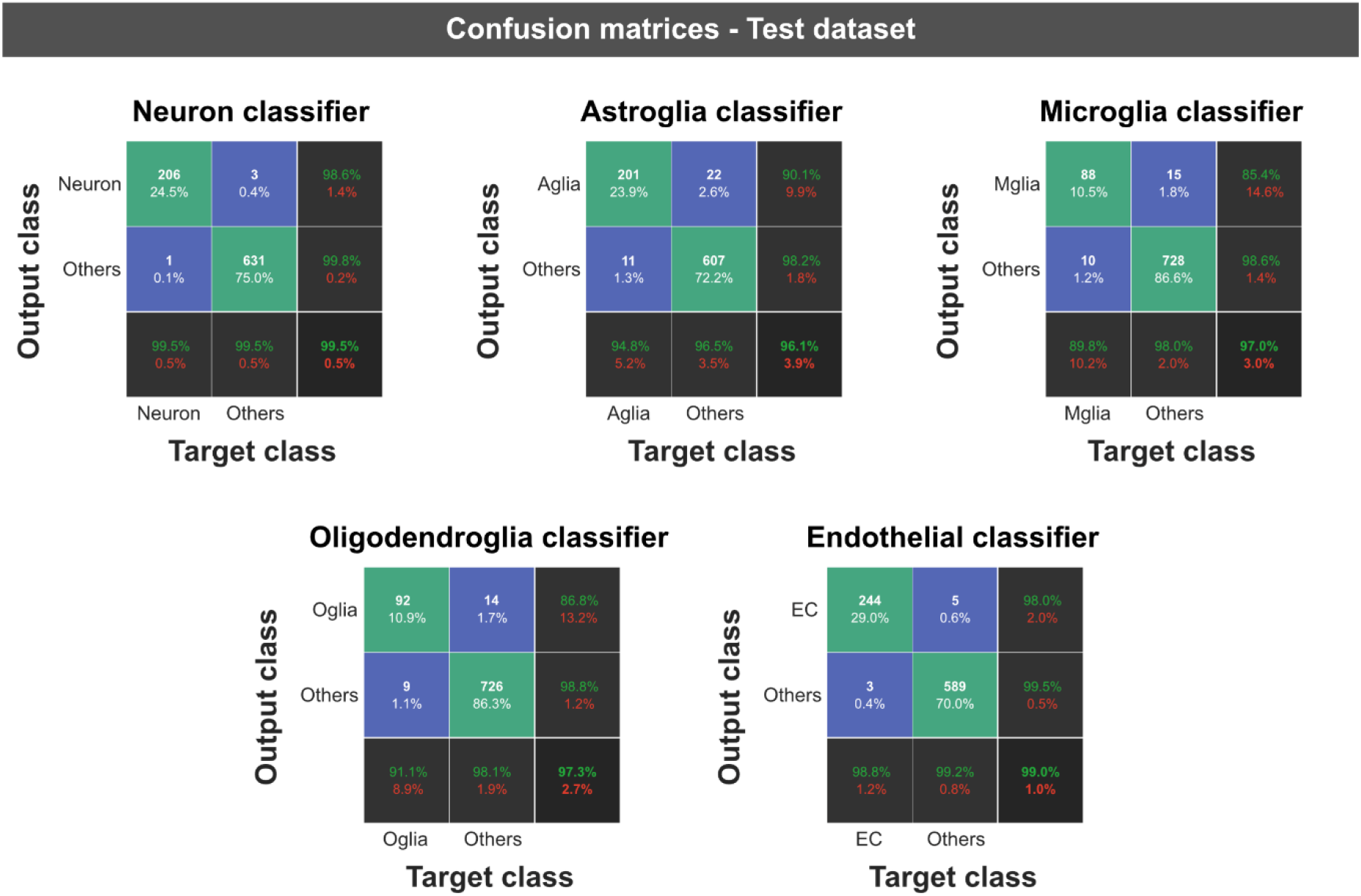
Confusion matrices for each classifier for the test dataset. Since each classifier distinguishes between the desired class and every “other” class, the confusion matrix consists only of 4 fields. Rows show the predicted class (Output Class), and the columns show the true class (Target Class). Green fields illustrate correct identification of target and “other” class, blue fields illustrate erroneous identifications. The number of observations and the percentage of observations compared to the total number of observations are shown in each cell. Column on the far right shows the precision (or positive predictive value) and false discovery rate in green and red respectively. Bottom row denotes recall (or true positive rate) and false negative rate in green and red. Cell on the bottom right shows overall accuracy of the classifier.

**Supplementary Figure 6.**
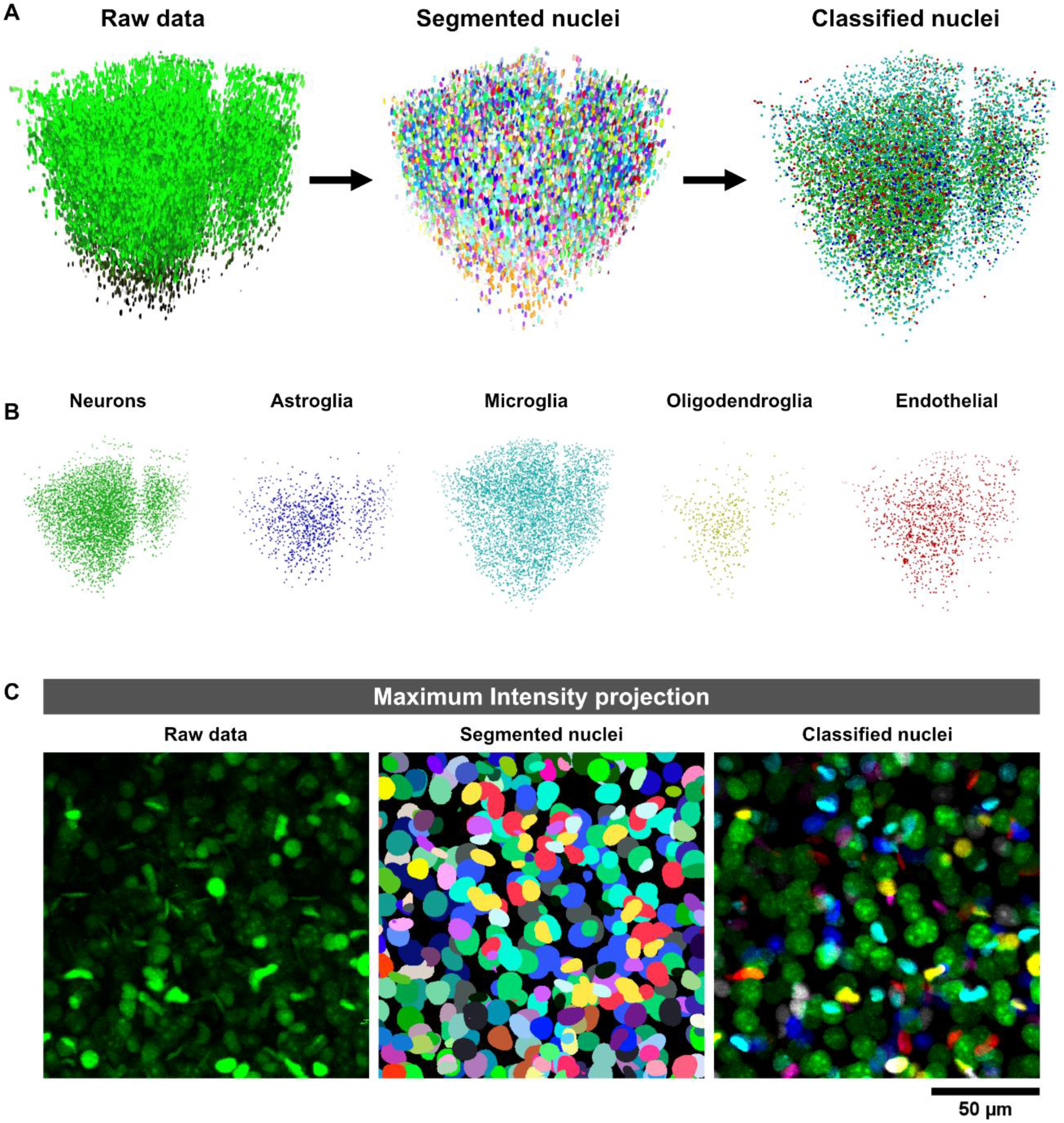
**(A)** 3-dimensional representation of raw, segmented and classified data (700 µm x 700 µm x 700 µm volume) visualizing comprehensive cell type composition. **(B)** Cell type distribution in a single volumetric stack. **(C)** Maximum intensity z-projection of a sub volume (150 µm x 150 µm x 100 µm) showing raw, segmented and classified nuclei in the X-Y axis.

**Supplementary Table 1:**
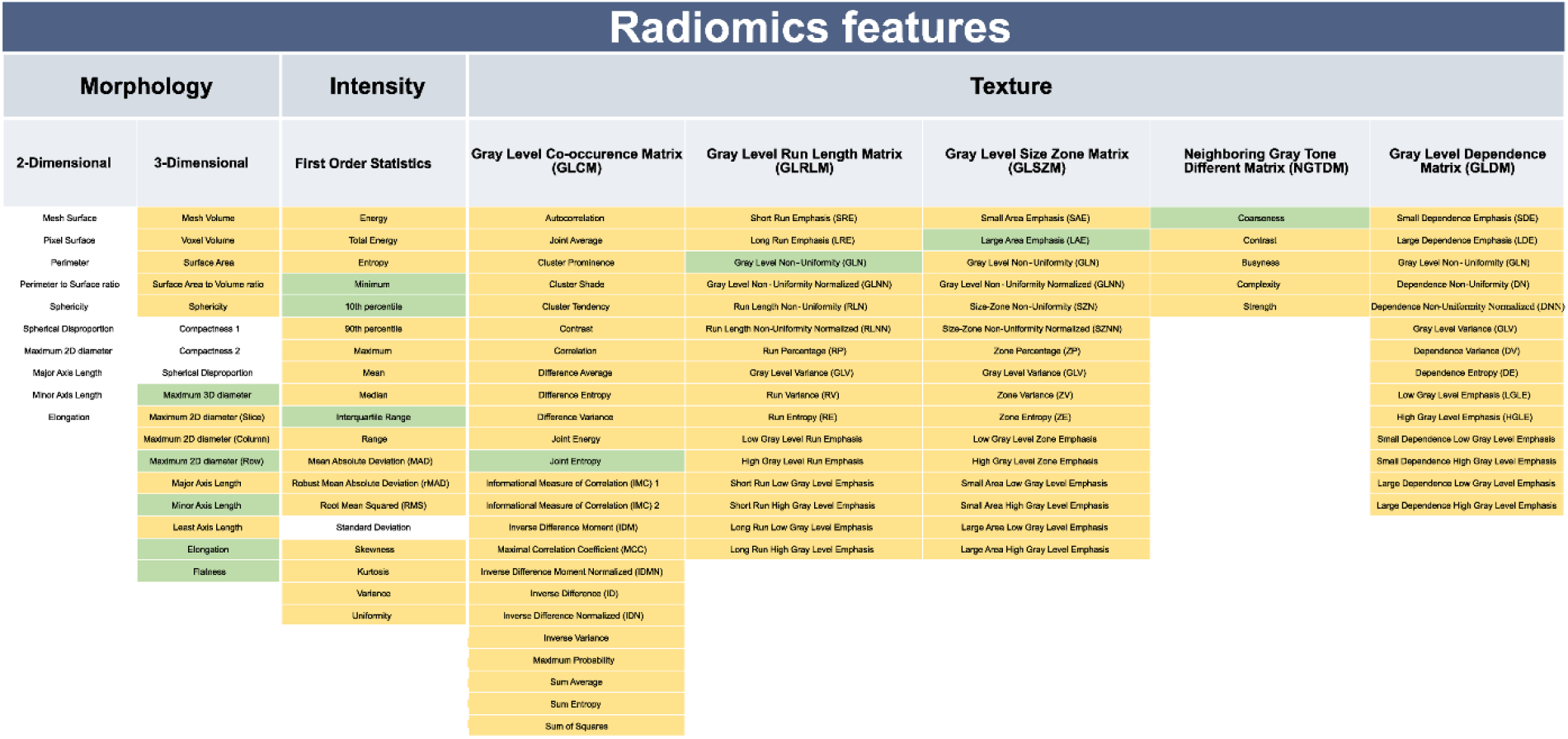
Radiomics features extracted from segmented nuclei. Features marked in green were used for cell type training and classification.

**Supplementary Table 2:**
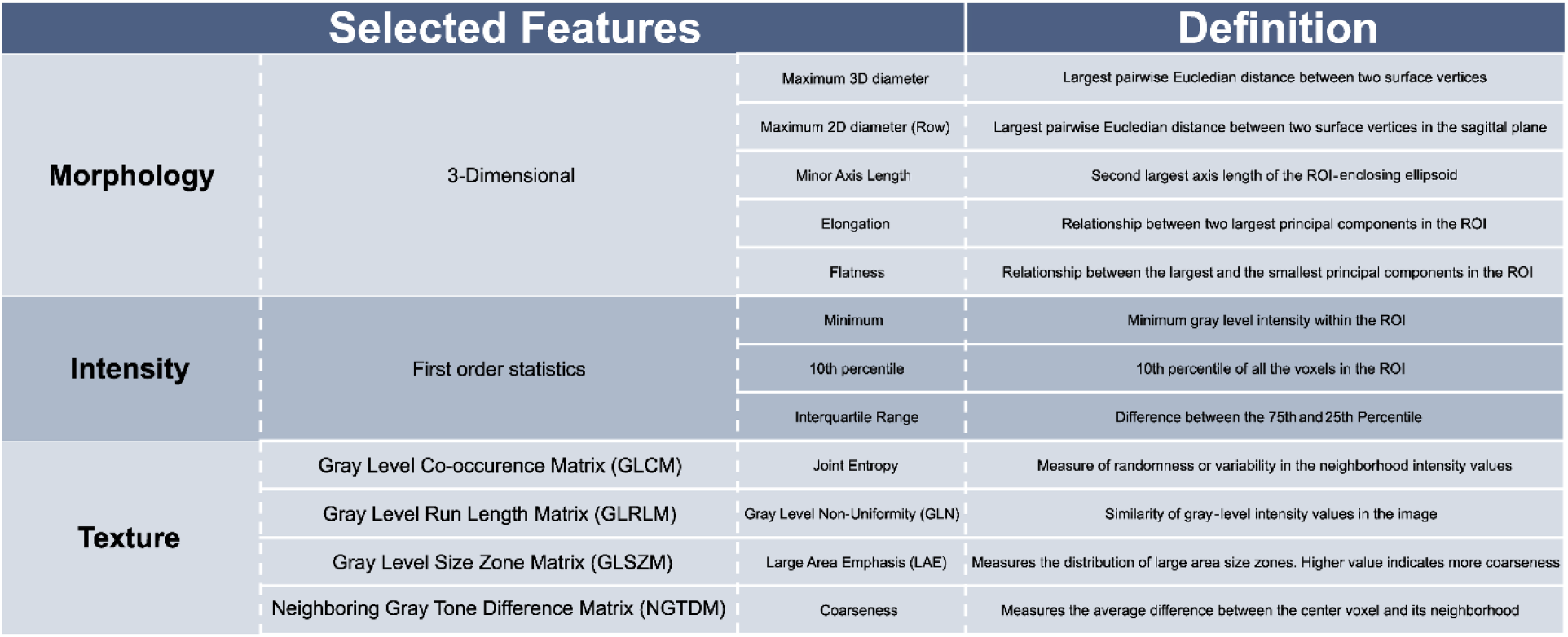
Morphology, intensity and texture features used for training and classification.

**Supplementary Text 1:** Explanations for technical terms used in the manuscript

Explanations for technical terms used in the manuscript:

**Accuracy:** Accuracy is a performance metric that quantifies the proportion of correctly classified data (both true positives and true negatives) out of all the data.

**Augmentation:** Augmentation involves applying various transformations, such as rotation, flipping, scaling, or adding noise, to the training data to increase its diversity. It helps improve the generalization and robustness of machine learning models.

**Binary mask**: A binary mask is a binary image where pixels are classified as either “foreground” (usually denoted as white) or “background” (usually denoted as black). Binary masks are commonly used to represent segmented regions, where each pixel is either part of the segmented structure (foreground) or not (background).

**Coarseness:** Coarseness is a texture feature that describes the scale of intensity variations within an image. It quantifies the contrast between regions of different intensities and their spatial frequency. A higher value indicates a lower spatial change rate and a locally more uniform texture.

**Confusion plots:** Confusion plots, also known as confusion matrices, are used to visualize the performance of a classification algorithm. They display the number of true positive, true negative, false positive, and false negative predictions made by the algorithm.

**Crops:** Crops refer to smaller regions or patches extracted from larger images.

**Entropy**: Entropy is a measure of the randomness or uncertainty in an image. It can be used to characterize the complexity of structures or textures within the image.

**Epoch:** An epoch is one complete cycle through the entire training dataset during the training process of a machine learning model in bioimage analysis.

**False discovery rate:** The false discovery rate (FDR) is a measure used to control the rate of false positive findings. It represents the proportion of incorrect positive predictions (false positives) among all positive predictions made by an algorithm.

**False positive / false negative:** False positive indicates an incorrect positive prediction, such as when the algorithm wrongly identifies background regions as part of the object of interest. False negative, on the other hand, refers to an incorrect negative prediction, where the algorithm fails to identify actual regions of interest.

**First order statistics:** First order statistics refers to the basic statistical measures computed directly from the pixel intensity values in an image, such as mean, median, standard deviation, etc.

**Flatness:** Flatness shows the relationship between the largest and smallest principal components in the region of interest (ROI) shape. The principal component analysis is performed using the physical coordinates of the voxel centers defining the ROI.

**Gray level co-occurrence matrix:** The gray level co-occurrence Matrix (GLCM) is a texture analysis method that quantifies the frequency of intensity value pairs at various spatial relationships within an image. It is commonly used to extract texture features.

**Gray level dependence matrix:** The gray level dependence matrix (GLDM) is a texture analysis technique that captures dependencies between pairs of pixels with specific intensity values in an image.

**Gray level run length matrix:** The gray level run length matrix (GLRLM) is a texture analysis technique that characterizes the lengths and frequencies of consecutive pixel runs with the same intensity value in an image.

**Gray level size zone matrix:** The gray level size zone matrix (GLSZM) is a texture analysis method that provides information about the distribution of connected regions of similar intensity values in an image.

**Gray level non-uniformity:** Gray level non-uniformity (GLNU) is a texture feature that quantifies the variance in pixel intensities within an image. It provides information about the spatial variations of grey levels in the image.

**Ground truth**: Ground truth refers to the manual or expert annotations of the correct regions of interest in an image. It serves as the gold standard for evaluating the performance of algorithms and machine learning models.

**Neighboring gray tone difference Matrix**: The neighboring gray tone difference matrix (NGTDM) is a texture analysis method that quantifies differences in pixel intensity values between neighboring pixels in an image.

**Overfitting:** Overfitting is a common problem in machine learning in general, where a model becomes too specialized on the training data and fails to generalize well to new, unseen data. It occurs when a model learns noise or irrelevant patterns from the training set.

**Precision:** Precision is a measure of the accuracy of positive predictions made by an algorithm. It calculates the proportion of true positive instances out of all positive predictions (true positives plus false positives).

**Radiomics:** Radiomics is a field in bioimage analysis that focuses on the extraction and analysis of quantitative features from images like texture, shape and morphology.

**Recall:** Recall, also known as sensitivity or true positive rate, is a measure of the algorithm’s ability to correctly identify positive instances (true positives) out of all actual positive instances in the image.

**Segmentation:** Segmentation refers to the process of identifying and delineating regions of interest (ROI) within an image, separating them from the background or other structures. The goal is to partition the image into meaningful regions that correspond to specific biological structures or objects of interest.

**Training:** Training involves using a dataset with known ground truth annotations to teach a machine learning model or algorithm how to perform the task, such as segmentation or classification, on new, unseen images.

**True positive / true negative:** True positive refers to correctly identified positive instances (e.g., correctly identified regions of interest). True negative, on the other hand, refers to correctly identified negative instances (e.g., background regions correctly identified as not being part of the object of interest).

**Validation:** Validation is the process of assessing the performance and generalization capabilities of a trained algorithm on a separate dataset. It helps to ensure that the model’s performance is consistent and reliable on new, unseen data.

